# The local topological free energy of the SARS-CoV-2 Spike protein

**DOI:** 10.1101/2021.02.06.430094

**Authors:** Quenisha Baldwin, Bobby G Sumpter, Eleni Panagiotou

## Abstract

The novel coronavirus SARS-CoV-2 infects human cells using a mechanism that involves binding and structural rearrangement of its spike protein. Understanding protein rearrangement and identifying specific residues where mutations affect protein rearrangement has attracted a lot of attention for drug development. We use a mathematical method introduced in [9] to associate a local topological/geometrical free energy along the SARS-CoV-2 spike protein backbone. Our results show that the total local topological free energy of the SARS-CoV-2 spike protein monotonically decreases from pre-to post-fusion and that its distribution along the protein domains is related to their activity in protein rearrangement. By using density functional theory (DFT) calculations with inclusion of solvent effects, we show that high local topological free energy conformations are unstable compared to those of low topological free energy. By comparing to experimental data, we find that the high local topological free energy conformations in the spike protein are associated with mutations which have the largest experimentally observed effect to protein rearrangement.

## 1 Introduction

The severe acute respiratory syndrome coronavirus 2 (SARS-CoV-2) has led to the COVID-19 global pandemic which has taken over 2 million lives. The need to stop the spread of the highly infectious virus requires disruption in the infection process and has become the focus for many scientists. Even more so, the task to be able to control and stop a global pandemic in the future is of great interest. Fusion of the membranes of both a host cell and a viral cell is necessary for infection [10]. Viral glycoproteins aid in this process by facilitating the binding of the two cells. Viral glycoproteins are folded proteins on the enveloped viral cell membrane which, when triggered, undergo irreversible dramatic conformational changes [14,19,24,42,48,62,63]. The SARS-CoV-2 S protein is a class I viral glyco-protein that consists in two subdomains (S1 and S2) and is triggered by cleavage at the S1 cleavage site [14,14,20,25,30,37,63]. S1, containing a receptor binding domain (RBD), binds to a host cell receptor, angiotensin-converting enzyme 2 (ACE2), and leads to a second cleavage at an S2’ cleavage site, adjacent to the fusion peptide. The protein undergoes several structural rearrangements that lead to a stable post-fusion state which brings the two membranes together [14]. In this manuscript, we quantify these changes with the aim to understand how local changes can trigger global conformations.

Folded proteins are defined by their primary, secondary, tertiary and quarternary structure [1]. The primary structure refers to the protein by amino acid sequence. The secondary structure refers to a sequence of 3-dimensional building blocks the protein attains (beta sheets, α-helices, coils). The tertiary structure refers to the 3-dimensional conformation of the entire polypeptide chain. The rearrangement of viral proteins during protein fusion changes both their tertiary structure and their secondary structure. We use tools from knot theory, namely, the Writhe and the Torsion, to characterize the viral protein conformations at the length-scale of 4 consecutive residues along the backbone.

In the last decades, measures from knot theory have been applied to biopolymers [3–8, 12,13,16,21,29,35,38,39,46,50,51,54,56,58] and in particular to proteins to classify their conformations [6–8,15,23,31,41,46,53,55]. One of the simplest measures of conformational complexity of proteins that does not require an approximation of the protein by a knot dates back to Gauss; the Writhe of a curve. In [9] the Writhe and the Torsion were used to define a novel topological/geometrical free energy that can be assigned locally to the protein. The results therein showed that high local topological free energy conformations are independent of the local sequence and may be involved in the rate limiting step in protein folding.

In this paper we apply this method to the spike protein of SARS-CoV-2 to characterize its conformation in various phases of viral fusion. Our results show that the local topological free energy of the spike protein is decreasing monotonically as the protein undergoes various conformational changes pre-fusion to post-fusion, in agreement with a transition of the protein from a metastable to stable state. Our results in combination with DFT calculations suggest that local conformations of high local topological free energy are unstable. By comparing our results to experimental data, we find that residues in high local topological free energy conformations are possible candidates for mutations with impact on protein rearrangement.

The paper is organized as follows: Section 2 describes the topological and geometrical functions for characterizing 3-dimensional conformation used in this paper and the density function theory calculations used to evaluate conformational stability. Section 3 describes the results of this method for SARS-CoV-2. Finally, in Section 4, we summarize the findings of our analysis.

## 2 Methods

In this Section we give some definitions necessary for the rest of the manuscript.

### 2.1 Measures of topological/geometrical complexity

We represent proteins by their CA atoms, as linear polygonal curves in space. A measure of conformational complexity of curves in 3-space is the Gauss linking integral. When applied to one curve, this integral is called the Writhe of a curve:

#### Definition 2.1.

(Writhe). For an oriented curve *l* with arc-length parameterization *γ*(*t*), the Writhe, *Wr*, is the double integral over *l*:

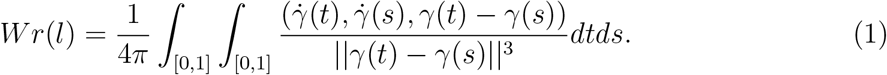

It is a measure of the number of times a chain winds around itself and can have both positive and negative values.

The total Torsion of the chain, describes how much it deviates from being planar and is defined as:

#### Definition 2.2.

The *Torsion* of an oriented curve *l* with arc-length parameterization γ(t) is the double integral over *l*:

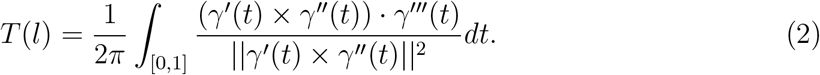

The Writhe and the Torsion have successfully been applied to study entanglement in biopolymers and proteins in particular [2,6,6–8,17,18,40,43–45,47,49,52].

An important property of the Gauss linking integral and the Torsion which makes them useful in practice is that they can be applied to polygonal curves of any length to characterize 3-dimensional conformations at different length scales. In this work, we use the Writhe and the Torsion to characterize the local conformation of parts of the protein at the length scale of 4 residues, we call this the *local Writhe (local Torsion*, resp.).

#### Definition 2.3.

We define the *local Writhe of a residue* (resp. *local Torsion*), represented by the CA atom *i* to be the Writhe (resp. *Torsion*) of the protein backbone connecting the CA atoms *i,i* + 1, *i* + 2, *i* + 3.

The local Writhe is a measure of the local orientation of a polygonal curve and a measure of its compactness. For example a very tight right handed turn (resp. left-handed) will have a positive (resp. negative) Writhe value close to 1 (resp. −1), while a relatively straight segment will have a value close to 0. Similarly, the Torsion is 0 for a planar segment and increases to ±1 as the segment deviates from being planar.

### 2.2 Topological/Geometrical free energy

In this Section we give the definition of the local topological free energy, originally defined in [9]. To assign a topological/geometrical free energy along a protein backbone, we first derive the distributions of the local Writhe and local Torsion and local ACN in the ensemble of folded proteins. To do this in practice, we use a curated subset of the crystal structures provided in the PDB [11]. Namely, we use the dataset of unbiased, high-quality 3-dimensional structures with less than 60% homology identity from [60]. Then for each residue of a given protein we compare its local Writhe (resp. Torsion) value to those of the ensemble and a free energy is assigned to the residue based on the population of that value in the ensemble.

Let *X* denote a topological parameter (local Writhe, local Torsion). Let *d_X_* denote the density (ie. the number of occurrences) of *X* in the folded ensemble (*d_Wr_,d_T_*, respectively). Let *m_X_* (resp. *m_T_*) denote the maximum occurrence value for *X*. To any value *p* of *X*, we associate a purely topological/geometrical free energy:

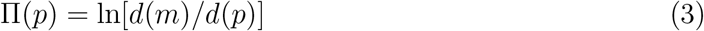

The total local topological free energy of a protein is defined as the sum of local topological free energies in Writhe (resp. Torsion) along the protein backbone. In [9] it was shown that the experimentally observed folding rates of a set of 2-state single domain proteins decrease with increasing total local topological free energy of the proteins.

We will say that a residue is *rare* or in high local topological free energy in a parameter *X* (local Writhe or local Torsion) if its value *X* = *p* is such that Π(p) ≥ *w*, where *w* is a threshold corresponding to the 95th percentile of Π-values across the set of folded proteins. We stress that our definition of rare residue involves 4 consecutive residues, starting from the one we label as rare. We will say that a residue is in a high local topological free energy conformation, we denote LTE, when it is one of the 4 residues composing a conformation with high local topological free energy. The 95^th^ percentile of the Π_Wr_-value distribution in the PDB culled set corresponds approximately to absolute Writhe values greater than 0.1. The 95^th^ percentile of the Π_T_-value distribution in the PDB corresponds approximately to absolute Torsion values greater than 0.3 or smaller than 0.1. Thus, high LTE in Writhe values correspond to high Writhe in absolute value, while high LTE in Torsion values may be values close to 0 in absolute value. This suggests that high LTE conformations in Writhe may not necessarily be high LTE in Torsion conformations and the opposite. In [9] it was found that high local topological free energy conformations are independent of local sequence. By comparison with experimental data it was shown that the number of such local conformations in a protein may be related to the rate limiting step in protein folding [9].

### 2.3 Density Function Theory (DFT) Calculations

Geometry optimizations of the identified high local topological free energy within the SARS-CoV-2 spike protein, were carried out in a model aqueous solution phase without imposing geometrical restrictions by using the NWChem suite of programs (version 7.0.2) [59]. All residues were optimized at the MO6-2X/6-311++G** level, e.g., via hybrid metafunctionals [66], and solvent effects were accounted for by using the Solvation Model Based on Density (SMD), [36]. The optimization of the identified residues via DFT was done to evaluate the validity of our hypothesis that high local topological free energy conformations are indicative of unstable structures.

## 3 Results

### 3.1 Local topological free energy of SARS-CoV-2

#### 3.1.1 Local topological free energy of viral glycoproteins pre- to post-fusion

The normalized (by its length) total local topological free energy of a set of representative examples of viral proteins of 3 main classes pre- and post-fusion is shown in Figure 2. We find that the total local topological free energy in Torsion decreases from pre-fusion to post-fusion for all proteins. Similarly, we find that the total local topological free energy in Writhe also decreases pre-fusion to post-fusion, with the exception of SARS. This suggests that viral proteins during fusion are guided towards a minimum of the total local topological free energy, in agreement with a transition from a metastable to a stable state.

**Figure 1:**
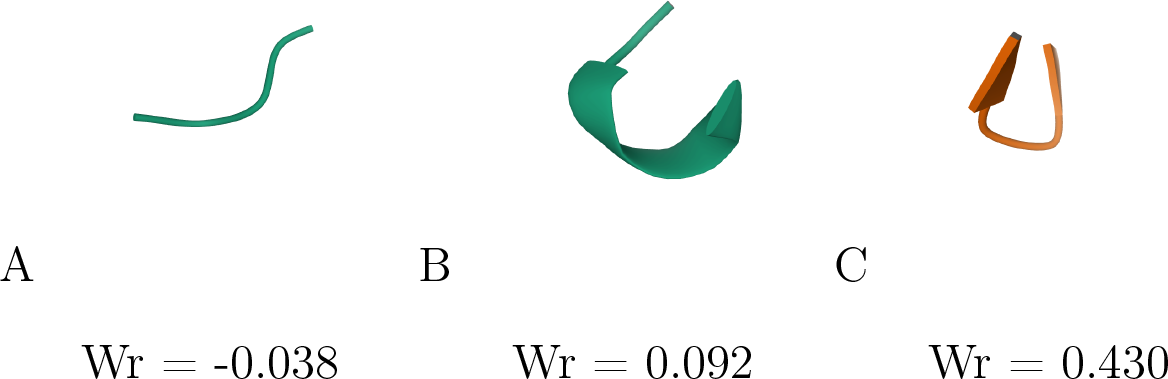
Example of Writhe values. (A) 16PK residues 1-4 (B) residues 4-8 and (C) 1GK9 residues 92-96 shown.

**Figure 2:**
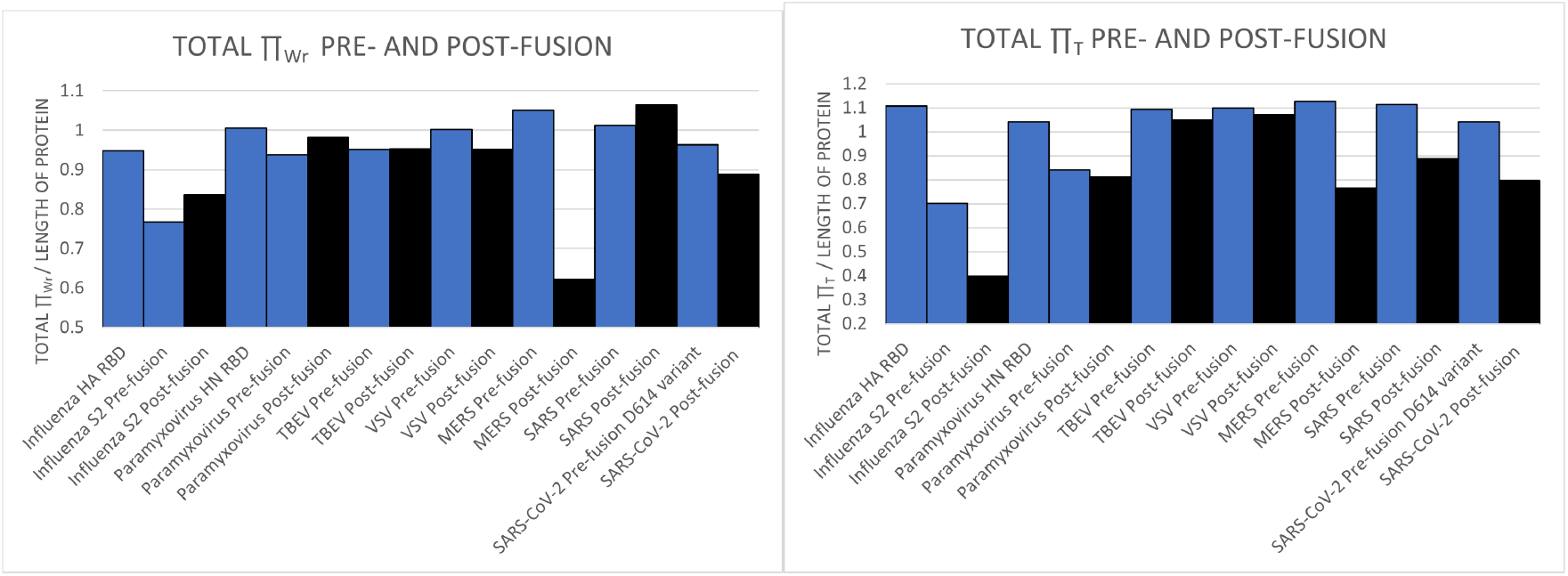
The normalized (by the length of the proteins) total local topological free energy in Writhe (Left) and Torsion (Right) of viral glycoproteins pre- (in blue) and post-fusion (in black). Some pre-fusion proteins are composed by two chains (one of which is the RBD) and are shown in two separate blue columns.

#### 3.1.2 Local topological free energy of SARS-CoV-2 from closed to open conformation

We found that the total topological free energy of SARS-CoV-2 decreases from pre- to post-fusion (see Figure 2). We next analyze how the local topological free energy changes in various pre-fusion stages. In pre-fusion the SARS-CoV-2 glycoprotein may be in uncleaved closed, cleaved closed, cleaved open and intermediate state (see Figure 8 Top). A closed conformation entails that all three RBD are in the down position [14]. An open conformation entails that there is an RBD in the up position, accessible for the angiotensin-converting enzyme 2 or ACE2 receptor to bind [14,30,64]. A cleaved protein indicates that the protein has been proteolytically cleaved at the cleavage site by a furin protease into the receptor binding subunit of S1. This is necessary for conformational changing of the RBD in human coronavirus. The fusion subunit of S2 remains associated after cleavage until post-fusion [65]. An intermediate conformation indicates that cleavage at the RBD has occurred and the RBD has been removed yet refolding has not occurred [14].

Figure 8 (Bottom Left) shows the total local topological free energy of the Spike protein of SARS-CoV-2 in Writhe in various stages of rearrangement pre-fusion. We see a continuous decrease of the local topological free energy from the closed uncleaved, closed cleaved, open and intermediate conformation in Writhe. We also find a decrease from the closed uncleaved to the intermediate conformation in Torsion (see SI).

Figure 8 (Bottom Right) shows the distribution of the total local topological free energy in each domain of SARS-CoV-2 normalized by the length of the domain in closed uncleaved, closed cleaved, open and intermediate conformations. Our results suggest that a decrease of the local topological free energy in a domain is associated with a rearrangement of the domain in 3-space and possibly with its involvement in protein rearrangement. Namely, we see that the total local topological free energy of the RBD, which is involved in the initial stage of pre-fusion rearrangement, decreases continuously. We see that the fusion peptide has overall much higher total local topological free energy in Writhe in the closed conformations, where it is protected, and decreases sharply in the open conformations, where it is exposed. From closed to open HR1 and S2, which are involved in pre-fusion rearrangement, also show a decrease. On the other hand, the CTD, which is involved in the protein rearrangement that follows after opening and binding from pre-fusion to post-fusion, shows an increase in local topological free energy. Similarly, we see an increase of the total local topological free energy of CH from the open to intermediate state. CH may play a role post-fusion in extending the protein.

#### 3.1.3 Local topological free energy of SARS-CoV-2 in D614, G614 and HexaPro mutations

In the previous sections we analyzed the structure of the SARS-CoV-2 D614 variant. This variant contains the S-2P stabilizing mutation. This is one of the most studied variants in the literature for which open/closed/intermediate crystal structures are reported. In this Section we compare this variant to two other variants of interest, G614 and HexaPro [28]. Our results in Figure 4 (Right), show that the G614 variant has an overall much higher local topological free energy in comparison to the HexaPro variant and the D614 variant. This is in agreement with experimental results which suggest that the D614 and HexaPro are more stable. The D614 variant is proposed to form a hydrogen bond with T859 (of the neighboring protomer or chain), limiting its flexibility while the G614 variant does not form this bond [34]. The HexaPro variant consists in 6 mutations [28] and has shown to stabilize the pre-fusion structure and produced high yield. In terms of the distribution of the local topological free energy in domains, we find that the G614 variant has increased local topological free energy in SD1, SD2, CD and HR1 domains. The results for Π_T_ are similar and are shown in SI.

**Figure 3:**
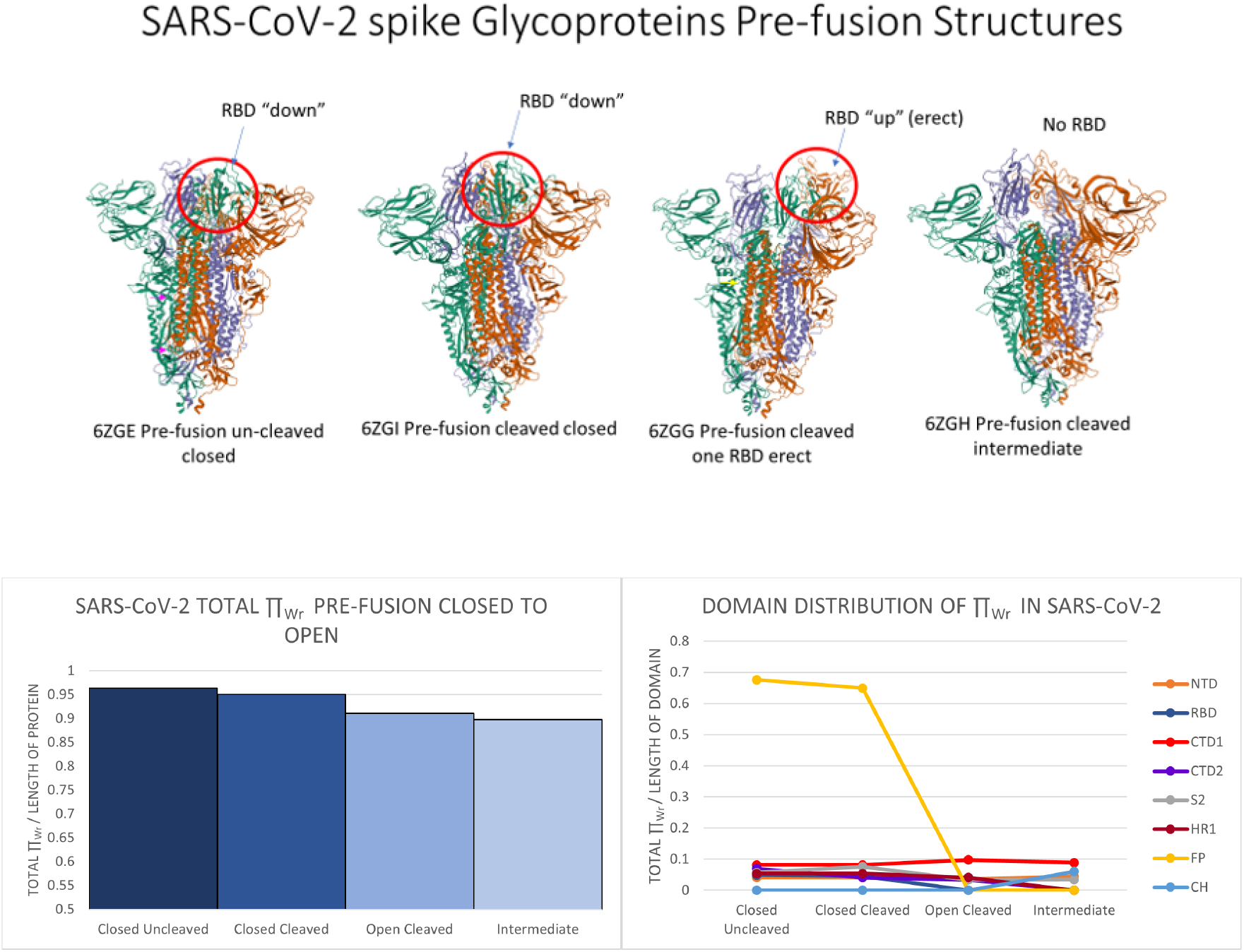
Top: From left to right, snapshots of SARS-CoV-2 pre-fusion protein at four stages: uncleaved closed (6ZGE), cleaved closed (6ZGI), cleaved open (6ZGG) and intermediate (6ZGH). The RBD is circled in red. Bottom Left: The normalized total local Π_Wr_ for SARS-CoV-2 protein at the 4 pre-fusion stages. Bottom Right: The normalized total local ⊓_wr_-values for SARS-CoV-2 protein domains at the 4 pre-fusion stages. Crystal structure images were pulled from the Protein Data Bank [11].

**Figure 4:**
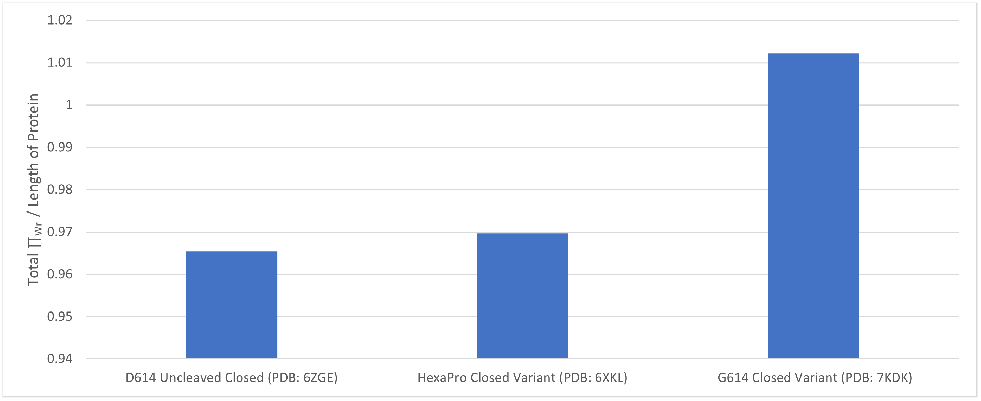
The normalized total Π_Wr_ for SARS-CoV-2 D614, G614 and HexaPro closed pre-fusion variants.

### 3.2 High local topological free energy conformations

In this section we focus on those local conformations in SARS-CoV-2 with high local topological free energy.

#### 3.2.1 DFT minimized local conformations

In this section we use DFT calculations to compare the minimal energy configurations of high local topological free energy versus medium/low topological free energy conformations in SARS-CoV-2.

We calculate the difference (Π*_Wr_*)*_DFT_* — (⊓*_Wr_*)*_PDB_*, between the Π*_Wr_* values of the DFT obtained conformations versus those of the PDB. We do this for the high local topological free energy conformations versus medium or low local topological free energy conformations in SARS-CoV-2. The distribution of the differences is shown in Figure 5. Orange indicates the difference for high local topological free energy conformations and blue for medium/low local topological free energy conformations.

**Figure 5:**
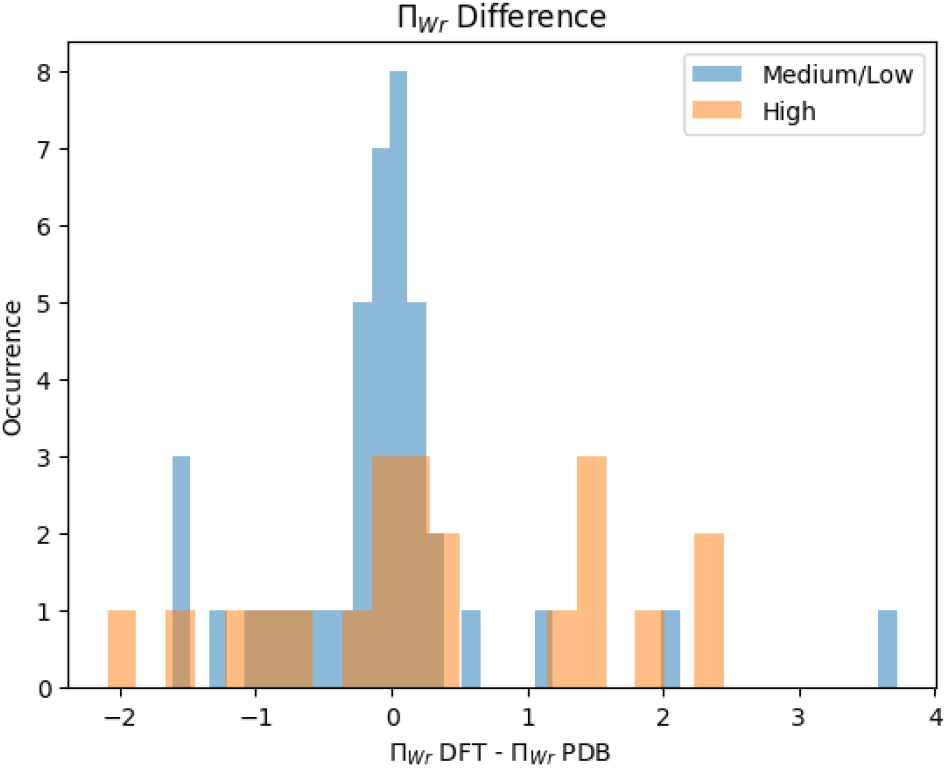
The distribution of (Π_Wr_)_DFT_ — (⊓_Wr_)_PDB_ The difference for residues in high local topological free energy conformations is shown in orange and that of medium or local topological free energy conformations is shown in blue. The distribution of high LTE minimized differences has a skewness of —0.006 while that of medium/low LTE has skewness of 1.717. We note that 2 of the largest positive differences in medium/low LTE (outliers in blue) are for conformations that proceed a gap in the PDB sequence.

Our results show that the distribution of (⊓*_Wr_*)*_DFT_* — (⊓*_Wr_*)*_PDB_* is more broad for the high LTE conformations compared to the medium/low LTE conformations. The skewness of the distribution of the medium/low LTE conformations is 1.717, while that of the high LTE conformations is —0.006. In particular, we find that for the majority of medium/low topological free energy conformations, their DFT structure is either very similar or it has lower local topological free energy. In contrast, many of the DFT reduced conformations of the high local topological free energy conformations, have even higher local topological free energy. These results further corroborate the idea that high local topological free energy conformations are indicative of unstable structures.

#### 3.2.2 High local topological free energy conformations in SARS-CoV-2

The high LTE conformations in the SARS-CoV-2 S protein at various stages pre-fusion, post-fusion and for some of its variants are given in Table 1 in SI. The local conformations are denoted by the first residue they are composed by. Thus the table lists only those initial residues.

**Table 1:**
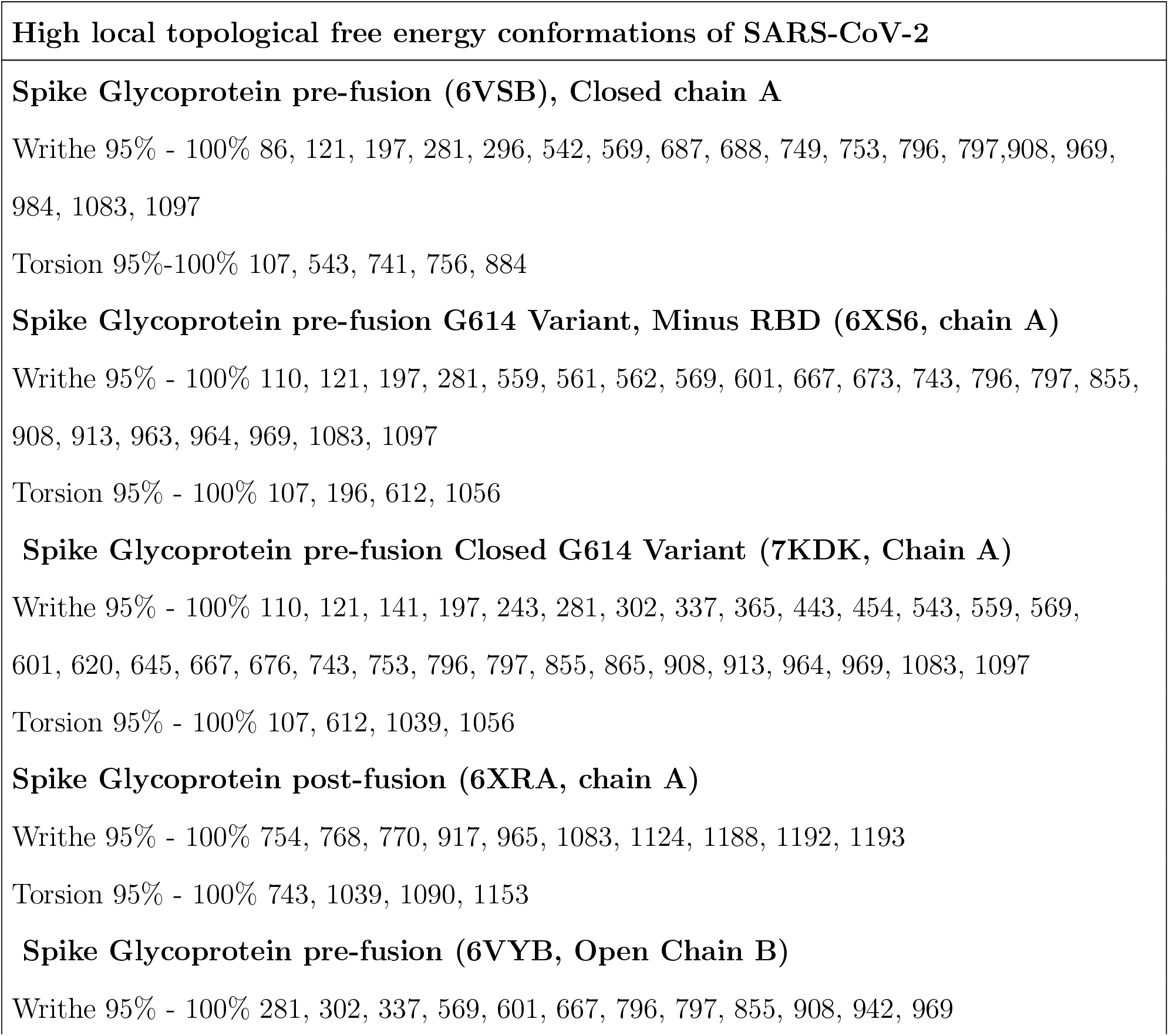
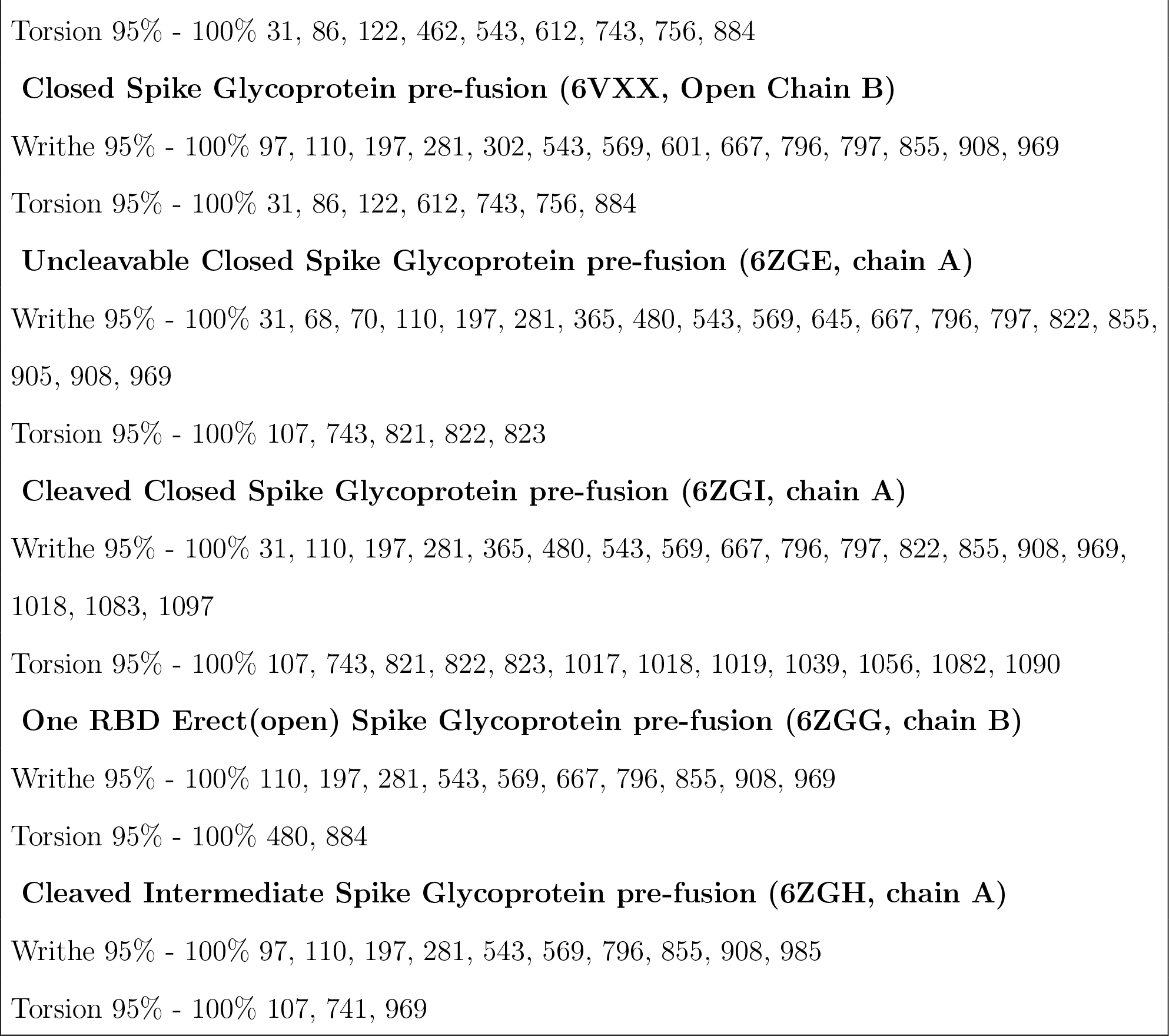
Residues in SARS-CoV-2 where High local topological free energy conformations of 4 residues begin. These correspond to the 95th-100th percentiles of the Writhe and Torsion distributions of the PDB culled ensemble.

The distribution of high local topological conformations in Writhe in the SARS-CoV-2 D614 and G614 variants pre-fusion domains and D614 post-fusion is shown in Figure 6. We find a higher number of high LTE in Writhe conformations in HR1 in the G614 variant than in D614. HR2 has a high number of high LTE conformations only post-fusion.

**Figure 6:**
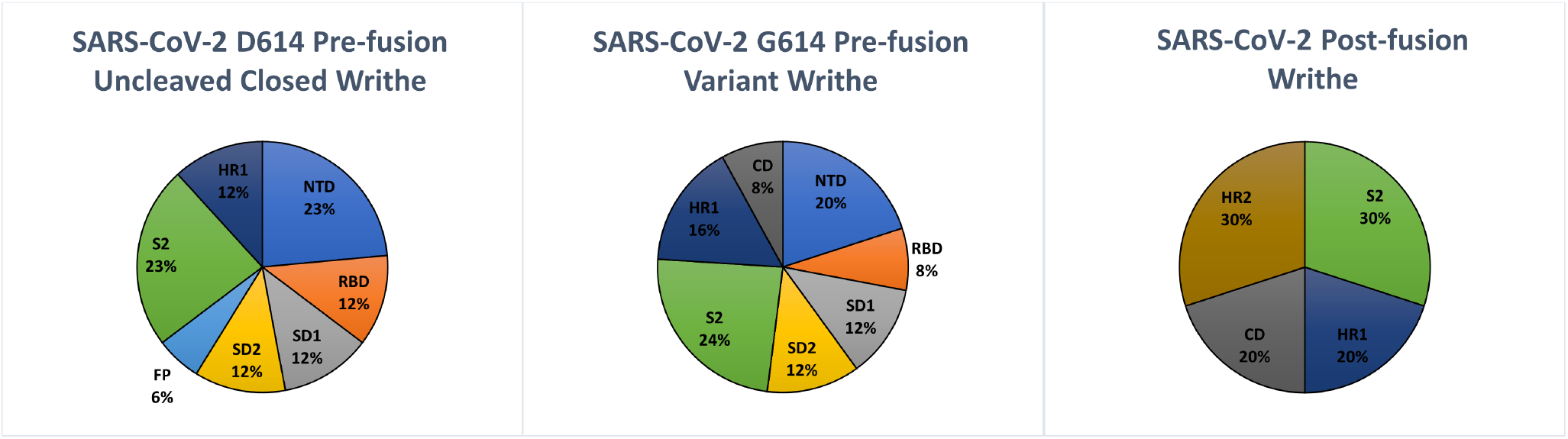
The distribution of high LTE conformations in domains in SARS-CoV-2 D614 and G164 variants pre-fusion and D614 post-fusion.

The number of high local topological free energy conformations in Writhe in the D614 variant in closed uncleaved, closed cleaved, open and intermediate states pre-fusion are shown in Figure 7. Overall, we see the same trends as those reported for the local topological free energy per domain, suggesting that the presence of high local topological free energy conformations signals rearrangement of a domain to a conformation with less high LTE conformations.

**Figure 7:**
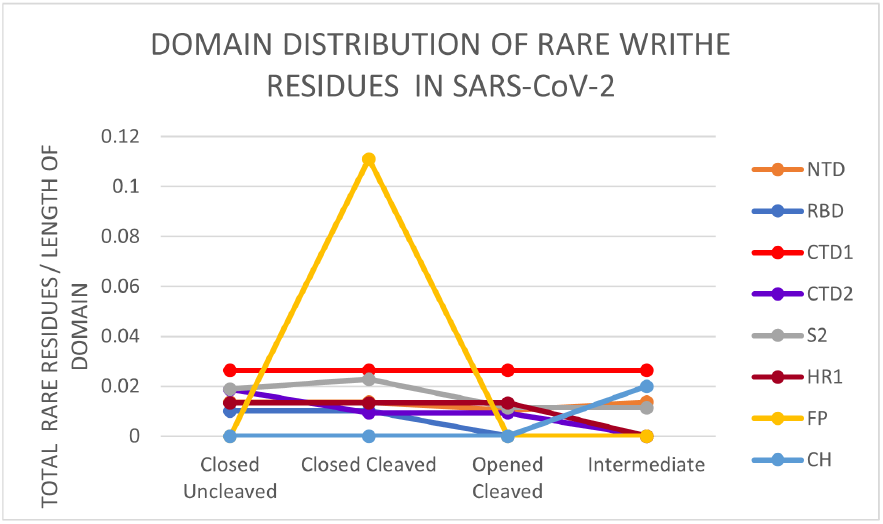
Distribution of high LTE conformations in SARS-CoV-2 pre-fusion domains in four stages: uncleaved closed (6ZGE), cleaved closed (6ZGI), one RBD erect (6ZGG) and cleaved intermediate (6ZGH).

**Figure 8:**
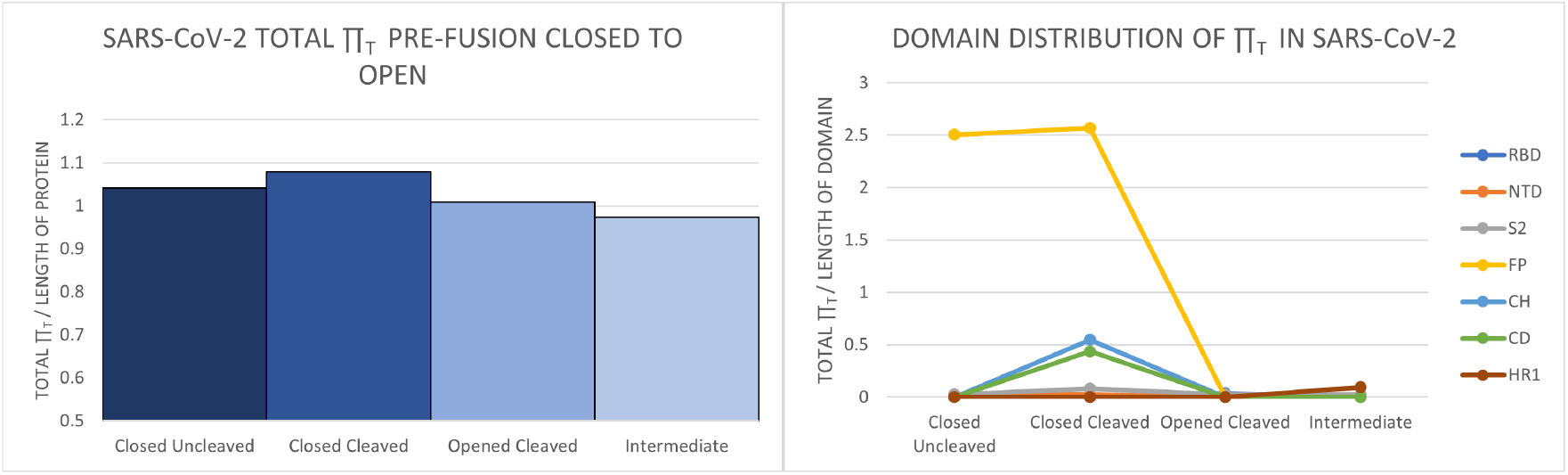
Left: The normalized local ⊓ for SARS-CoV-2 pre-fusion proteins in motion in Torsion. Right: The normalized local ⊓-values for SARS-CoV-2 pre-fusion domains in Torsion.

**Figure 9:**
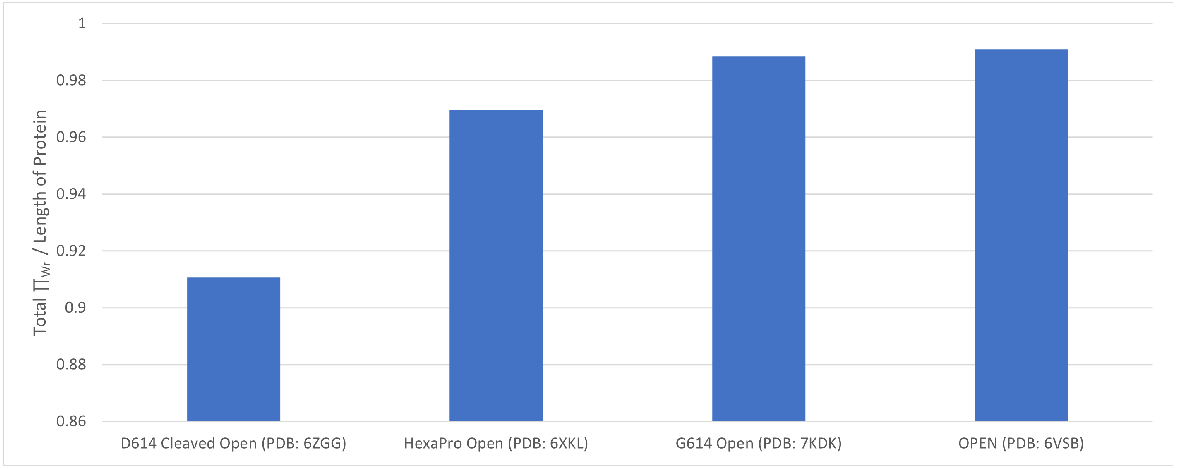
The normalized total Π_Wr_ for SARS-CoV-2 cleaved open (1 RBD erect) D614, G614 open and HexaPro open pre-fusion variants. Also shown here is the normalized total Π_Wr_ of 6VSB, a D614 variant that consists only of the 2P mutation.

#### 3.2.3 Mutations and high local topological free energy

In this section we compare the experimentally reported ability of a mutation to impact 3-dimensional rearrangement and the local topological free energy at that site for SARS and SARS-CoV-2 known mutations. Our results are shown in Tables 2 and 3 in SI. In the following all high LTE conformations are in Writhe, unless stated otherwise.

**Table 2:**
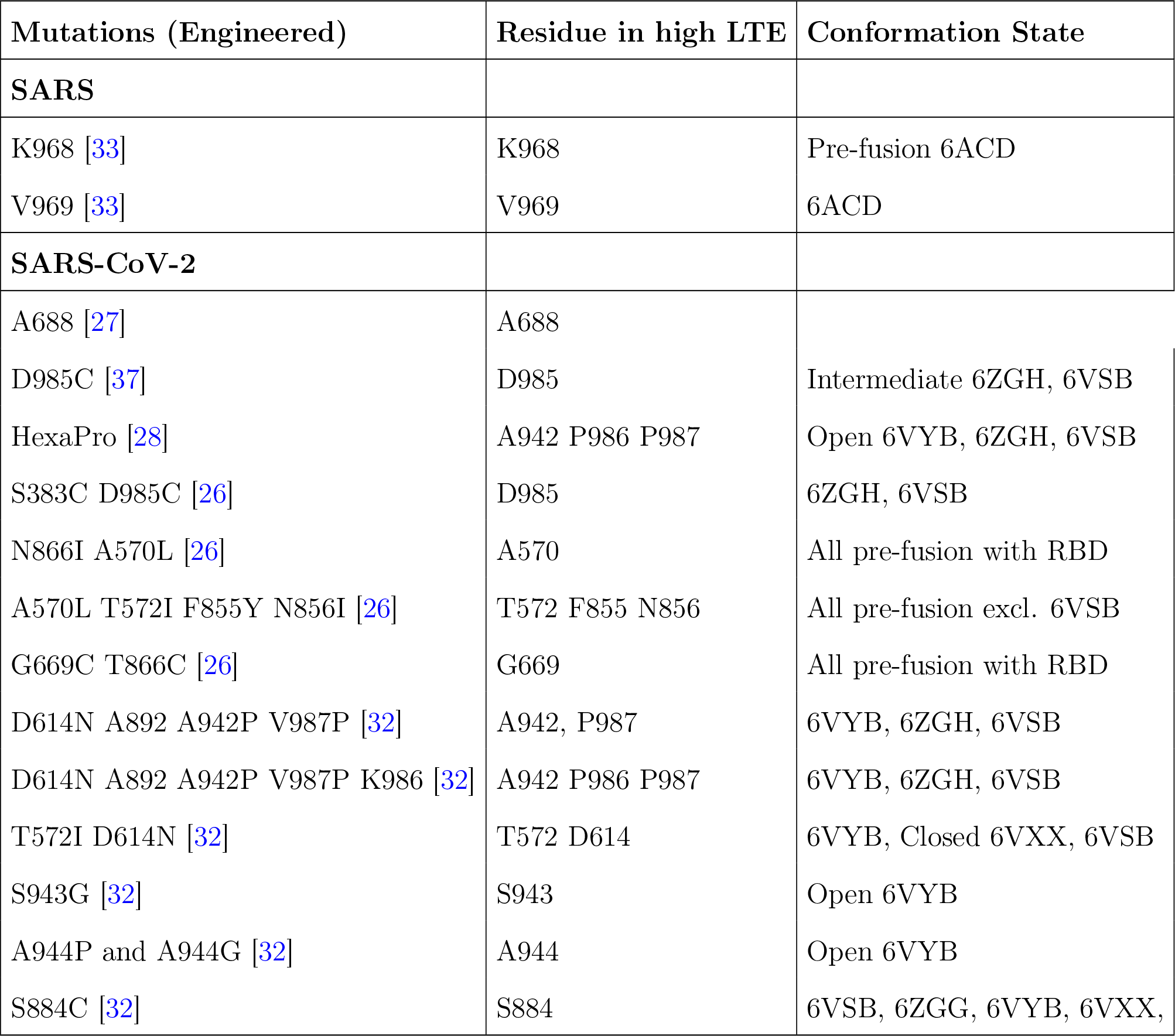
Mutations of SARS-CoV-2 spike protein with experimentally observed impact on protein rearrangement.

**Table 3:**
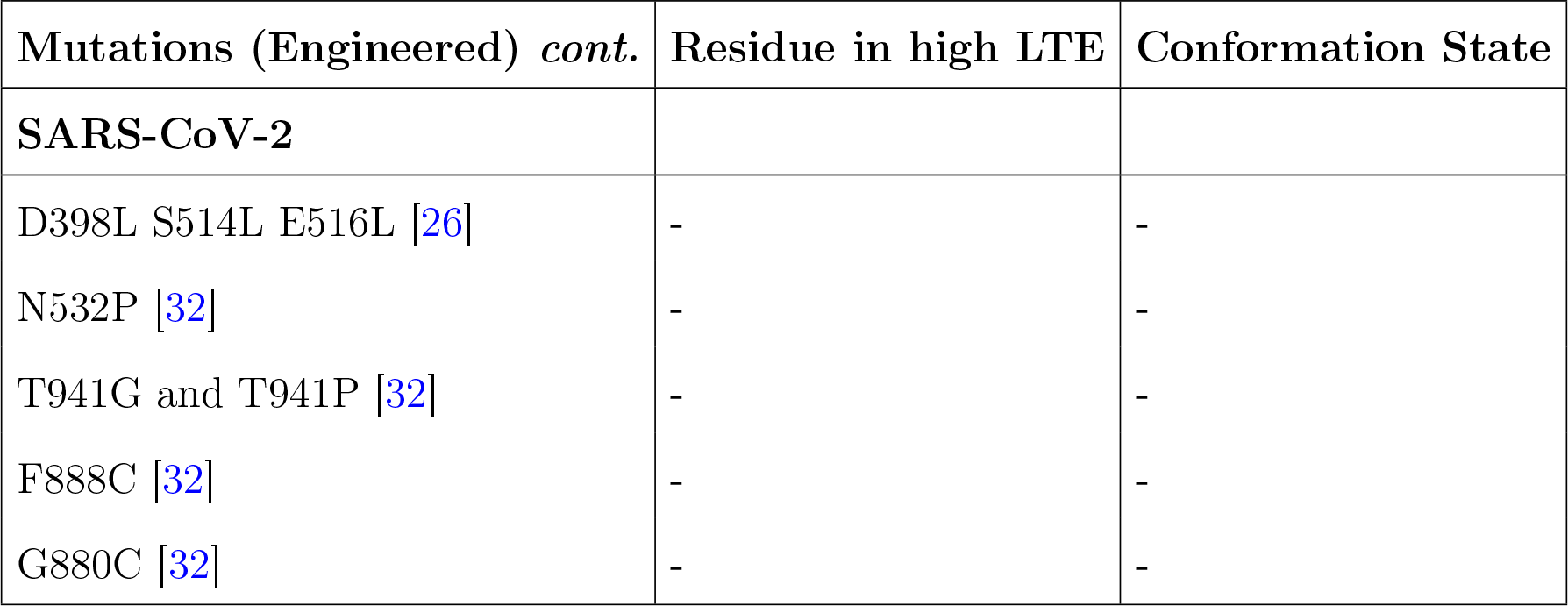
Continuation of Table 2. These are mutations that where not found in high LTE conformations.

| Mutations (Engineered) *cont.* | Residue in high LTE | Conformation State |
| --- | --- | --- |
| **SARS-CoV-2** |  |  |
| D398L S514L E516L [26] | - | - |
| N532P [32] | - | - |
| T941G and T941P [32] | - | - |
| F888C [32] | - | - |
| G880C [32] | - | - |

Overall, we find that 75% of the mutations which are known experimentally to change the 3-dimensional properties of the spike protein are at residues in high local topological free energy conformations. The effect on protein conformation of the new mutants that have naturally arisen is to our knowledge undetermined. We find that only 42% of those natural mutants occurred at residues in high local topological free energy conformations. However, we note that the only one of the natural occurring mutants for which we have a crystal structure is G614. We found that the residue 614 was in a high LTE in Torsion conformation in D614, only in the open state, while it is found in a high LTE in Torsion conformation in the closed mutant G614.

More precisely, mutations at SARS residues K968 and V969 (known as 2P a double proline mutation) caused disruption of conformational changes upon binding [33]. We find both residues in high local topological free energy conformations.

The cleavage site of SARS-CoV-2, residue A688, which has been identified as important in viral rearrangement [27], is in a high local topological free energy conformation. The residue 614 where the G614 natural mutation occurred is in a high local topological free energy conformation in Torsion for D614 and G614 [14]. Also, residue 985 which is involved in a stabilizing mutation, is in a high local topological free energy conformation [37]. In [28] 43 substitutions where studied. The most efficient was the HexaPro variant (involves 6 mutations), which stabilized the pre-fusion structure and produced high yield [28]. We find two out of the six mutations composing HexaPro (F817P, A892P, A899P, A942P, K986P, V987P) to be in a high local topological free energy conformation before the mutation and none to be in a high local topological free energy conformation after the mutation.

In [26], using a different method, specific sites for mutation that would change global conformation were identified. These were a double cysteine mutant, S383C D985C (RBD to S2 double mutant (rS2d)), a triple mutant, D398L S514L E516L (RBD to NTD (triple mutant (rNt)), a double mutant, N866I A570L (subdomain 1 to S2 double mutant (u1S2d)), a quadruple mutant, A570L T572I F855Y N856I (subdomain 1 to S2 quadruple mutant (u1S2q)) and finally, a double cysteine mutant, G669C and T866C, to link SD2 to S2 (subdomain 2 to S2 double mutant (u2S2d)). We found that out of these 5 mutants, 4 contained residues in high local topological free energy conformations. Moreover, one of the most efficient mutations contained 2 residues in a high local topological free energy conformation.

In [32], a combination of mutations were examined. Categorized by location the mutations included N532P, T572I, D614N, D614G of SD1, and A942P, T941G, T941P, S943G, A944P, A944G, and A892P, F888 and G880C, S884C and A893C, and K986P and V987P of the C-terminus of HR1 [32]. We find residues 572, 614, 942, 943, 944, 884, 986, 987 are in high local topological free energy conformations (884 in Torsion). A892P was reported to increase closed trimers while A942 decreased closed trimers. The double proline mutation (2P) at K986P and V987P stabilized the pre-fusion structure. K986P was reported to have higher ACE2 binding affinity while D614N and T572I reported to have low binding affinity [32].

The top 10 naturally prevalent mutations of the SARS-CoV-2 S protein (the recently discovered UK and South African mutations [22,57]) are shown in Table 4. These mutations are are believed to increase infectivity and transmission and the majority of them are located in S1 [61]. The precise impact of these mutations is not yet clear but current data suggests that these mutations may increase both transmission and virulence of the virus. Both the UK and South African variants are believed to increase the binding affinity because some are located within the receptor binding motif of the RBD, residues which directly bind to the ACE2 receptor. The UK variant is reported to contain several mutations including deletion mutations (at residues 69, 70, 144) and substitution mutations (at 501, 570, 614, 681, 716, 982, 1118), shown in Table 4. One of these residues, residue 570, is in a high local topological free energy conformation. The South African variant, involves N501Y, E484K, and K417N. We find none of these residues to be in high local topological free energy conformations before the mutation.

**Table 4:**
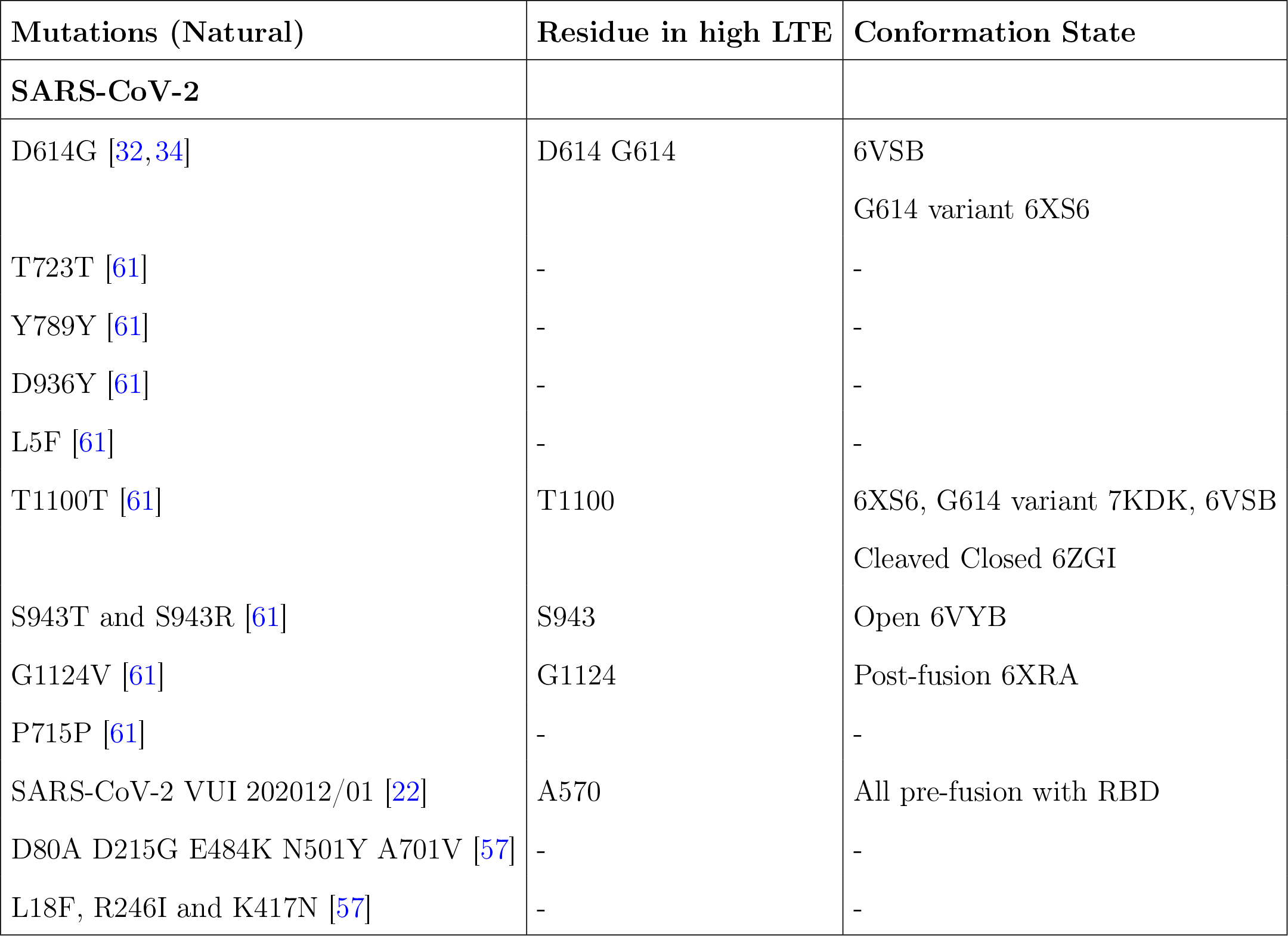
Mutations of emerging SARS-CoV-2 spike protein variants.

## 4 Discussion

We used the local topology/geometry of protein crystal structures alone to associate a local topological free energy to the protein backbone. We find that total local topological free energy decreases from pre- to post-fusion. In addition the total local topological free energy of the spike protein of SARS-CoV-2 pre-fusion decreases continuously in the steps leading to protein rearrangement, in agreement with a transition to an energetically more stable state. We found that the total local topological free energy of the G614 mutant was much higher than the D614 and HexaPro mutants. Experimental results have shown that the G614 mutant is more unstable compared to the D614 and HexaPro mutants. This finding further supports that the purely topological free energy can quantify protein stability.

The distribution of the local topological free energy in the domains of the spike protein changes from closed to open states and from pre- to post-fusion. Our results suggest that high total local topological free energy in domains indicates activity of the domain in protein rearrangement.

DFT calculations showed that local conformations of the spike protein of medium/low LTE relax to conformations with even lower LTE, while the conformations with high LTE relax to both lower and higher LTE conformations. This further supports that the high LTE conformations are more unstable. Comparing to experimental data of mutations known to enhance or obstruct protein rearrangement and stability of the Spike protein, we find that most of the mutations reported to alter protein conformation are at residues in high local topological free energy conformations.

## 5 Acknowledgments

QB and EP thank the support of NSF REU 1852042 and internal support of the University of Tennessee at Chattanooga. QB and EP thank the support of NSF DMS 1913180. BGS acknowledges work supported at Oak Ridge National Laboratory’s Center for Nanophase Materials Sciences, a US Department of Energy Office of Science User Facility.

## 6 Supplementary Information

### 6.1 Table of High local topological free energy residues

In this supplemental section, we report all the residues where a high local topological free energy conformation of 4 consecutive residues along a chain begins for SARS-CoV-2 pre-fusion, post-fusion and intermediate states as well as the G614 mutant.

### 6.2 Table of mutations

### 6.3 Domains of SARS-CoV-2 Spike protein

SARS-CoV-2 is a Class I fusion protein. It is composed of two subunits, the binding subunit (S1) and the fusion subunit (S2). Upon triggering by low pH or protein interaction or cleavage, the binding subunit is responsible for the binding of the virus to the host cell receptor which then allows for an irreversible structural rearrangement. The fusion subunit is responsible for facilitating fusion. The two subunits are connected by an *S_1_*/*S_2_* cleavage site.

S1 is composed by the receptor binding subunit, also called receptor binding domain (RBD) and sometimes also by the N-terminal domain (NTD) and the C-terminal domain (CTD). S1 dissociates from the rest of the viral protein from pre- to post-fusion. S2 is composed by a fusion peptide (FP) and a transmembrane domain (TMD) which both are used to insert into opposite membranes to join the membranes together. Other domains exist which we denote as domains with an associated number (we denote DN, where D is domain and N is a number). For SARS-CoV-2, additional domains in S2 include the heptad repeat region (HR1 and HR2), central helix (CH), cytoplasm domain (CD) and subdomain 3 (SD3). The HR and SD3 (post-fusion [20]) are domains located at the tail of the protein. HR1 rearranges during fusion in a fashion that plunges the FP into the host membrane. HR2, located at the tail of the protein, folds to bring the viral membrane to close proximity to the host membrane during fusion [30].

## References

[1] B Alberts, A. Johnson, J. Lewis, M. Raff, K. Roberts, and P. Walter. Molecular Biology of the Cell. New York: Garland Science, 2002.

[2] J. Arsuaga, T. Blackstone, Y. Diao, E. Karadayi, and M. Saito. The linking of uniform random polygons in confined spaces. J. Phys. A: Math. Theor., 40:1925–36, 2007.

[3] J. Arsuaga, Y. Diao, T. Kaplan, and M. Vazquez. The effects of density on the topological structure of the mitochondrial dna from trypanosomes. Journal of Mathematical Biology, 64:1087–1108, 2012.

[4] J. Arsuaga, M. Vazquez, P. McGuirk, S. Trigueros, D. W. Sumners, and J. Roca. Dna knots reveal a chiral organization of dna in phage capsids. Proc. Natl. Acad. Sci. (USA), 102:9165–9169, 2005.

[5] J. Arsuaga, M. Vazquez, S. Trigueros, D. W. Sumners, and J. Roca. Knotting probability of dna molecules confined in restricted volumes: Dna knotting in phage capsids. Proc. Natl. Acad. Sci. USA, 99:5373–5377, 2002.

[6] M. Baiesi, E. Orlandini, F. Seno, and A. Trovato. Exploring the correlation between the folding rates of proteins and the entanglement of their native state. J. Phys. A: Math. Theor., 50:504001, 2017.

[7] M. Baiesi, E. Orlandini, F. Seno, and A. Trovato. Sequence and structural patterns detected in entangled proteins reveal the importance of co-translational folding. Scientific Reports, 9:1–12, 2019.

[8] M. Baiesi, E. Orlandini, A. Trovato, and F. Seno. Linking in domain-swapped protein dimers. Scientific Reports, 6:1–11, 2016.

[9] Q. Baldwin and E. Panagiotou. The local topological free energy of proteins. bioRxiv 2021.01.06.425494 (submitted for peer-review), 2021.

[10] S. Belouzard, J. K. Millet, B. N. Licitra, and G. R. Whittaker. Mechanisms of coronavirus cell entry mediated by the viral spike protein. Viruses, 4:1011–1033, 2012.

[11] H. M. Berman, J. Westbrook, Z. Feng, G. Gilliland, T. N. Bhat, H. Weissig, I. N. Shindyalov, and P. E. Bourne. Magnetic helicity in a periodic domain. Nuc. Ac. Res., 28:235–242, 2000.

[12] D. Buck and Flapan. E. Predicting knot or catenane type of site-specific recombination products. J. Mol Biol., 374:1186–1199, 2007.

[13] D. Buck and Flapan. E. A topological characterization of knots and links arising from site-specific recombination. J. Phys. A.: Math. Theor., 40:12377–12395, 2007.

[14] Y. Cai, J. Zhang, T. Xiao, H Peng, R. M. Sterling, S. M. and Walsh Jr., S. Rawson, S. Rits-Volloch, and B. Chen. Distinct conformational states of sars-cov-2 spike protein. Science, 369(6511):1586–1592, 2020.

[15] P. Dabrowski-Tumanski, M. Piejko, S. Niewieczerzal, A. Stasiak, and J. I. Sulkowska. Protein knotting by active threading of nascent polypeptide chain exiting from the ribosome exit channel. J. Phys. Chem. B., 122:11616–11625, 2018.

[16] I. Darcy, J. Luecke, and M. Vazquez. Tangle analysis of difference topology experiments: applications to a mu protein-dna complex. Algebraic and Geometric Topology, 9:2247–2309, 2009.

[17] Y. Diao, A. Dobay, and A. Stasiak. The average inter-crossing number of equilateral random walks and polygons. J. Phys. A: Math. Gen., 38:7601–7616, 2005.

[18] Y. Diao, C. Ernst, K. Hinson, and U. Ziegler. The mean-squared writhe of alternating random knot diagrams. J. Phys. A: Math. Theor., 43:495202, 2010.

[19] L. J. Earp, S. E. Delos, H. E. Park, and J. M. White. The many mechanisms of viral membrane fusion proteins. 285:25–66, 2005.

[20] X. Fan, D. Cao, L. Kong, and X. Zhang. Cryo-em analysis of the post-fusion structure of the sars-cov spike glycoprotein. Nature Communications, 11:3618, 2020.

[21] E. Flapan, A. He, and H. Wong. Topological descriptions of protein folding. PNAS, 116:9360–9369, 2019.

[22] European Centre for Disease Prevention and Control. Rapid increase of a sars-cov-2 variant with multiple spike protein mutations observed in the united kingdom. ECDC, 2020.

[23] D. Goundaroulis, N. Gügümcu, S. Lambropoulou, J. Dorier, A. Stasiak, and L. H. Kauffman. Topological methods for open-knotted protein chains using the concepts of knotoids and bonded knotoids. Polymers, 9:444, 2017.

[24] S. C. Harrison. Mechanism of membrane fusion by viral envelope proteins. Advances in Virus Research, 64:64:231–361, 2005.

[25] S. C. Harrison. Viral membrane fusion. Virology, 0:498–507, 2015.

[26] R. et al. Henderson. Controlling the sars-cov-2 spike glycoprotein conformation. Nature Strut. Mol. Biol., 27:925–933, 2020.

[27] M. Hoffmann, H. Kleine-Wber, and S. Pohlmann. A multibasic cleavage site in the spike protein of sars-cov-2 is essential for infection of human lung cells. Molecular Cell, 78:1–6, 2020.

[28] C-L Hsieh and et al. Structure-based design of prefusion-stabilized sars-cov-2 spikes. Science, 369:1501–1505, 2020.

[29] X. Hua, B. Raghavan, D. Nguyen, J. Arsuaga, and M. Vazquez. Random state transitions of knots: a first step towards modeling unknotting by type ii topoisomerases. Topology and its applications, 157:1381–1397, 2007.

[30] Y. Huang, C. Yang, X. Xu, W. Xu, and S. Liu. Structural and functional properties of sars-cov-2 spike protein: potential antivirus drug development for covid-19. Acta Pharmacologica Sinica, 41:1141–1149, 2020.

[31] M. Jamroz, W. Niemyska, E. J. Rawdon, A. Stasiak, K. C. Millett, P. Sulkowski, and J. Sulkowska. Knotprot: a database of proteins with knots and slipknots. Nucleic Acids Res., 43:D306–14, 2015.

[32] J. Juraszek, L. Rutten, S. Blokland, and et al. Stabilizing the closed sars-cov-2 spike trimer. S. Nat Commun, 12:244, 2021.

[33] R. N. Kirchdoerfer, N Wang, J. Pallesen, D. Wrapp, H. L. Turner, C. A. Cottrell, K. S. Corbett, B. S. Graham, J. S. McLellan, and A. B. Ward. Stabilized coronavirus spikes are resistant to conformational changes induced by receptor recognition of proteolysis. Sci. Reports, 8:15701, 2018.

[34] B. et al. Korber. Tracking changes in sars-cov-2 spike: Evidence that d614g increases infectivity of the covid-19 virus. Cell, 2020.

[35] D. Marenduzo, E. Orlandini, A. Stasiak, D. W. Sumners, L. Tubiana, and C. Micheletti. Dna-dna interactions in bacteriophage capsids are responsible for the observed dna knotting. PNAS, 106:22269–22274, 2009.

[36] A. V. Marenich, C. J. Cramer, and D. G. Truhlar. Universal solvation model based on solute electron density and on a continuum model of the solvent defined by the bulk dielectric constant and atomic surface tensions. J. Phys. Chem., 113:6378–6396, 2009.

[37] M. McCallum, A. C. Walls, J. E. Bowen, D. Corti, and D. Veesler. Structure-guided covalent stabilization of coronavirus spike protein trimers in the closed conformation. Nat. Struct. Biol., 27:942–949, 2020.

[38] C. Micheletti, D. Marenduzzo, E. Orlandini, and D. W. Sumners. Knotting of random ring polymers in confined spaces. J. Chem. Phys., 124:64903.1–10, 2006.

[39] C. Micheletti and H. Orland. Efficient sampling of knotting-unknotting pathways for semiflexible gaussian chains. Polymers, 9:196, 2017.

[40] K. C. Millett, A. Dobay, and A. Stasiak. Linear random knots and their scaling behavior. Macromolecules, 38:601–606, 2005.

[41] W. Niemyskal, D. Dabrowski-Tumanski, M. Kadlof, E. Haglund, P. Sułkowski, and J. I. Sulkowska. Complex lasso: new entangled motifs in proteins. Scientific Reports, 6:36895, 2016.

[42] S. Nisole and A. Saïb. Early steps of retrovirus replicative cycle. Retrovirology, 1:9:1742–4690, 2004.

[43] E. Panagiotou and M. Kröger. Pulling-force-induced elongation and alignment effects on entanglement and knotting characteristics of linear polymers in a melt. Phys. Rev. E, 90:042602, 2014.

[44] E. Panagiotou, M. Kröger, and K. C. Millett. Writhe and mutual entanglement combine to give the entanglement length. Phys. Rev. E, 88:062604, 2013.

[45] E. Panagiotou, K. C. Millett, and P. J. Atzberger. Topological methods for polymeric materials: characterizing the relationship between polymer entanglement and viscoelasticity. Polymers, 11:11030437, 2019.

[46] E. Panagiotou and K. W. Plaxco. A topological study of protein folding kinetics. Topology and Geometry of Biopolymers, AMS Contemporary Mathematics Series, 746:223–233, 2020.

[47] E. Panagiotou, C. Tzoumanekas, S. Lambropoulou, K. C. Millett, and D. N. Theodorou. A study of the entanglement in systems with periodic boundary conditions. Progr. Theor. Phys. Suppl., 191:172–181, 2011.

[48] R. C. Penner. Backbone free energy estimator applied to viral glycoproteins. Journal of Computational Biology, 27:1–14, 2020.

[49] J. Portillo, Y. Diao, R. Scharein, J. Arsuaga, and M. Vazquez. On the mean and variance of the writhe of random polygons. J. Phys. A: Math. Theor., 44:275004, 2011.

[50] M. Pouokam, B. Cruz, S. Burgess, M. Segal, M. Vazquez, and J. Arsuaga. The rabl configuration limits topological entanglement of chromosomes in budding yeast. Scientific Reports, 9:6795, 2019.

[51] E. J. Rawdon, J. C. Kern, M. Piatek, P. Plunkett, A. Stasiak, and K. C. Millett. Effect of knotting on the shape of polymers. Macromolecules, 41:8281–8287, 2008.

[52] P. Rogen and B. Fain. Automatic classification of protein structure by using gauss integrals. Proc. Natl Acad. Sci, 100:119–24, 2003.

[53] K. Shimokawa, K. Ishihara, I. Grainge, D. Sherratt, and M. Vazquez. Ftsk-dependent xercd-dif recombination unlinks replication catenanes in a stepwise manner. PNAS, 110:20906–20911, 2013.

[54] R. Stolz, M. Yoshida, R. Brasher, M. Flanner, K. Ishihara, D. J. Sheratt, K. Shimokawa, and M. Vazquez. Pathways of dna unlinking: a story of stepwise simplification. Sci. Reports, 7:12420, 2017.

[55] J. I. Sulkowska, E. J. Rawdon, K. C. Millett, J. N. Onuchic, and A. Stasiak. Conservation of complex knotting and slipknotting in patterns in proteins. PNAS, 109:E1715, 2012.

[56] D. W. Sumners and S. G. Whittington. Untangling dna. Math Intelligencer, 12:71–80, 1990.

[57] H. Tegally and et al. Emergence and rapid spread of a new severe acute respiratory syndrome-related coronavirus 2 (sars-cov-2) lineage with multiple spike mutations in south africa. medRxiv, 2020.

[58] S. Trigueros, J. Arsuaga, M. E. Vazquez, D. W. Sumners, and J. Roca. Novel display of knotted dna molecules by two-dimensional gel electrophoresis. Nucleic Acids Research, 29:e67, 2001.

[59] M. Valiev, E.J. Bylaska, N. Govind, K. Kowalski, T.P. Straatsma, H.J.J. van Dam, D. Wang, J. Nieplocha, E. Apra, T.L. Windus, and W.A. de Jong. Nwchem: a comprehensive and scalable open-source solution for large scale molecular simulations. Comput. Phys. Commun., 181:1477, 2010.

[60] G. Wang and R. L. Jr Dunbrack. Pisces: a protein sequence culling server. Bioinformatics, 19:1589–1591, 2003.

[61] R. Wang, Y. Hozumi, C. Yin, and G. W. Wei. Decoding sars-cov-2 transmission and evolution and ramifications for covid-19 diagnosis, vaccine, and medicine. Journal of chemical information and modeling, 60(12):5853–5865, 2020.

[62] W. Weissenhorn, A. Hinz, and Y. Gaudin. Virus membrane fusion. FEBS Letters, 581:2150–2155, 2007.

[63] J. M. White, S. E. Delos, M. Brecher, and K. Schornberg. Structures and mechanisms of viral membrane fusion proteins: multiple variations on a common theme. PLOS Comput Biol, 43:189–219, 2008.

[64] D. Wrapp, N. Wang, K. S. Corbett, J. A. Goldsmith, C-L. Hsieh, O. Abiona, B. S. Graham, and J. S. McLellan. Cryo-em structure of the 2019-ncov spike in the prefusion conformation. Science, 367:1260–1263, 2020.

[65] A.G. Wrobel, D.J. Benton, and P. et al. Xu. Sars-cov-2 and bat ratg13 spike glycoprotein structures inform on virus evolution and furin-cleavage effects. Nat Struct and Mol Bio, 27:763–767, 2020.

[66] Y. Zhao and D. G. Truhlar. A new local density functional for main-group thermochemistry, transition metal bonding, thermochemical kinetics, and noncovalent interactions. J. Chem. Phys., 125:194101, 2006.

